# Soil nitrogen cycling rates are linked to microbial functional and taxonomic groups across the United States

**DOI:** 10.64898/2026.04.01.715970

**Authors:** Corinne R. Vietorisz, Chikae Tatsumi, Zoey R. Werbin, Jennifer M. Bhatnagar

**Affiliations:** Boston University, Department of Biology, Boston, MA, USA; Okinawa Institute of Science and Technology, Onna, Okinawa, Japan

## Abstract

Soil microbes support life on Earth by regulating the availability of nutrients in soils, yet we lack a fundamental, baseline knowledge of which fungi and bacteria are associated with specific soil nitrogen (N) cycling processes across ecosystems. We identified functional and taxonomic groups of fungi and bacteria that are associated with net ammonification and nitrification rates in soils from diverse ecosystems across the United States, including the environmental contexts where these relationships exist. To accomplish this, we co-analyzed soil, microbial, plant, and climatic data from 19 sites across the U.S. National Ecological Observatory Network (NEON). Distinct microbial groups were associated with net ammonification versus nitrification rates, highlighting the need to measure and model these two processes separately. The relative abundance of several microbial groups known for their N-decomposition abilities (i.e., Acidobacteriae, Bacteroidia, Saccharomycetes yeasts, ectomycorrhizal fungi) were positively associated with net ammonification rates across diverse environmental conditions. Meanwhile, pathogenic fungi, copiotrophic bacteria, and bacterial classes containing denitrifying bacteria were positively associated with net nitrification rates in many wet, hot, and high-N environments. These results deepen our understanding of soil microbiome ecology and represent a practical starting point to develop microbial-explicit biogeochemical cycling models at large spatial scales.

## INTRODUCTION

The transformation of elements through soils supports life on Earth by controlling nutrient availability for plants, microbes, and all organisms that depend upon them, including humans^1^. Complex microbial communities drive nutrient cycling by decomposing organic matter^2^ and transforming nutrients into new chemical forms^3^, including as plant-available nutrients or greenhouse gases^4^. Microbial groups are also impacted by nutrient cycling rates, as microbial community composition often shifts when nutrient availability changes^5^. Yet, we lack a fundamental, baseline knowledge of which fungi and bacteria are associated with specific nutrient cycling processes at large spatial scales and across diverse ecosystems. This is a major issue in environmental science, because it prevents us from including important biological controls over nutrient cycling in models that simulate and predict future global ecosystem functions, such as biogeochemical cycling, leaf litter decomposition, plant productivity, and greenhouse gas emissions^6^. Additionally, understanding the ties between microorganisms and nutrient cycling may aid in predicting the abundances of key microbial groups that affect plant and animal health or global climate, such as pathogens or greenhouse gas-producing denitrifying bacteria^7,8^.

The complexity of microbial communities has made it difficult to determine relationships between microbial taxa and specific nitrogen (N) cycling processes at large spatial scales (e.g., across hundreds of kilometers, as is simulated in Earth Systems Models^6^). Recently, it has been suggested that grouping microbial taxa into functional groups based on their ecological guild (a group of microbes that share similar ecological lifestyles and functions) or taxonomic group can serve as a starting point for organizing microbial diversity into meaningful categories that can be explicitly incorporated into biogeochemical models^6^. However, to our knowledge, the associations between microbial functional or taxonomic groups and N cycling rates have not been tested at large geographic scales–only at single sites or single ecosystem types, often temperate forests. Additionally, these relationships may be dependent on vegetation, climate, and soil factors due to different optimal environmental conditions for each microbial group. For example, in temperate forests, net N mineralization or ammonification rates (the release of ammonium from organic sources in soil) are positively correlated with the relative abundance of ectomycorrhizal fungi (EMF), which are mutualistic with tree roots and produce N-decomposing enzymes, as well as oligotrophic bacteria, which degrade N and thrive in low-carbon and low-nutrient environments^9–11^. However, EMF and oligotrophic bacteria are abundant in colder, N-limited systems^12–16^, such that these relationships might only hold in these environments. Additionally, there is debate around the ecological significance of functional groupings like the bacterial oligotroph/copiotroph categorizations, and it remains largely unknown whether these categorizations are helpful for explaining ecosystem processes like ammonification^14,16^. Meanwhile, in temperate forests, soil net nitrification rates (the conversion of ammonium to nitrate) can correlate positively with the relative abundance of copiotrophic (carbon and nutrient-loving) bacteria^10,17^ and ammonia-oxidizing microbes^9,18,19^, which perform the rate-limiting step in nitrification^20^. Copiotrophic bacteria are abundant in energy-rich ecosystems, including warmer, wetter, N-rich environments that favor nitrification^14–16^, which can also favor microbial groups such as pathogenic fungi^12,20^, so it is unclear which specific microbial taxa are related to nitrification under these conditions.

In most studies, ammonification and nitrification are binned together into a single rate of “net N mineralization” (net inorganic N release from breakdown of soil organic matter), even though ammonification and nitrification are performed by separate groups of microbes^20^. This, combined with the fact that microbial-N cycling relationships have mostly been explored within individual ecosystem types, makes it difficult to differentiate when and where microbial functional groups are meaningfully associated with ammonification versus nitrification processes.

In this study, we sought to determine how functional groups of fungi and bacteria are associated with net ammonification and nitrification rates across large geographic regions and the environmental context-dependencies of those associations. We hypothesized that EMF and oligotrophic bacteria would be positively associated with net ammonification rates in cold, N-limited ecosystems, while pathogenic fungi and copiotrophic bacteria would be positively associated with net nitrification rates in warmer, wetter, N-rich environments. To test this hypothesis, we quantified relationships between the relative abundances of fungal and bacterial functional and taxonomic groups and soil net ammonification and nitrification rates across 19 sites of the U.S. National Ecological Observatory Network (NEON), spanning the contiguous United States, Alaska, and Puerto Rico (Fig. 1a). The variation in soil net N mineralization and nitrification rates across NEON sites has been difficult to explain due to extreme environmental and spatial variation^22^, so we tested for microbial-N cycling relationships across the entire dataset, as well as within specific environmental conditions where they are hypothesized to occur (Supplementary Table 1) using a combination of non-linear correlation, multiple regression, and machine learning analyses. Because abiotic and vegetation variables within and across sites were often structured by latitude and longitude (Fig. 1b), our analyses accounted for spatial autocorrelation in microbial communities. We found strong relationships between N cycling processes and specific microbial functional and taxonomic groups, as well as key environmental thresholds for the presence or absence of these relationships.

**Fig. 1.**
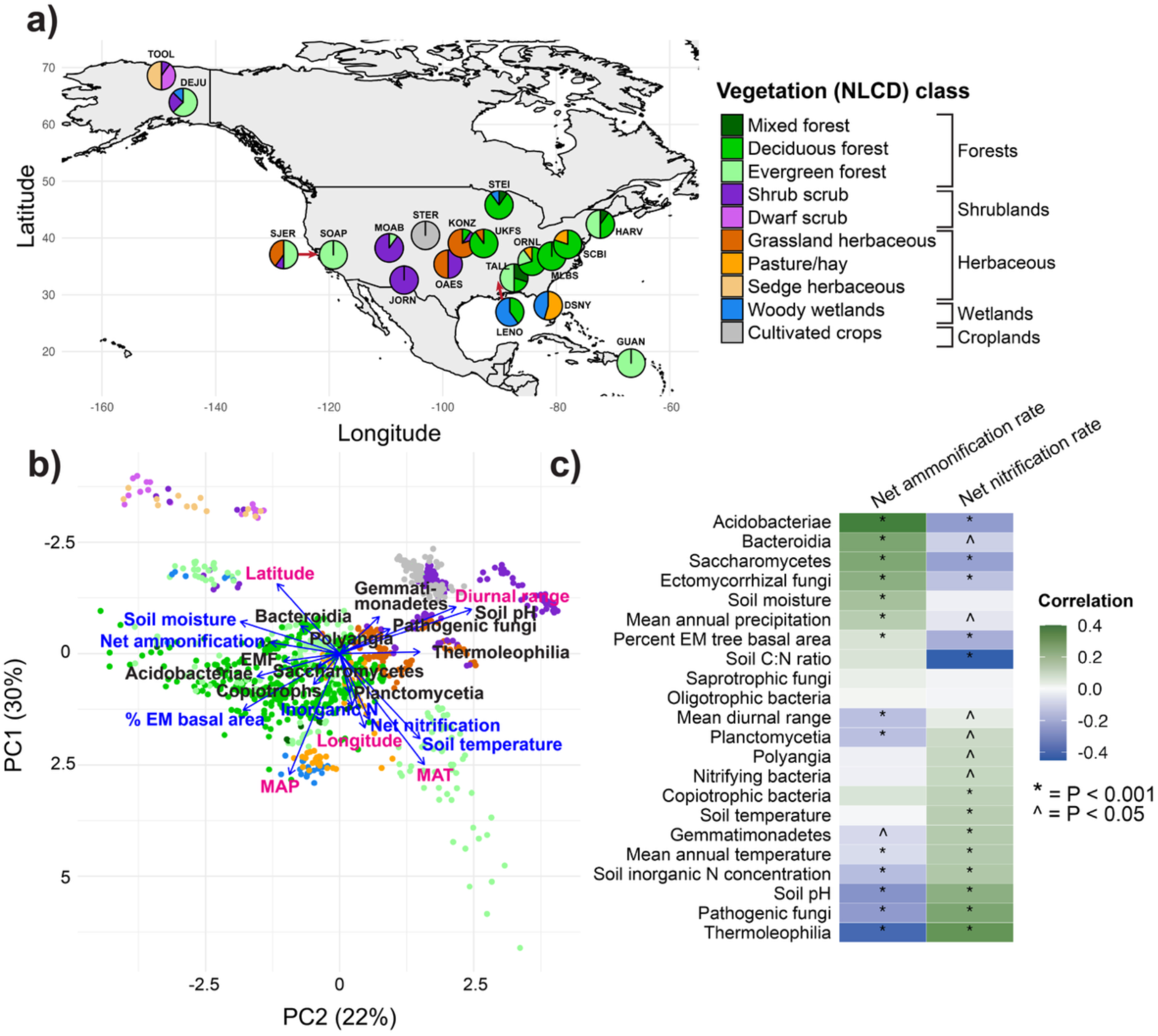
Study system and correlates with net ammonification and nitrification rates across the NEON data set. a) NEON sampling sites used in this study, where each pie chart represents one sampling site and colors within pie charts represent the proportion of plots belonging to a vegetation cover class (according to the National Land Cover Database, NLCD). All sites contain 8 - 10 plots. Site codes are shown adjacent to each pie chart and correspond to the site codes in Supplementary Table 5. Charts for the sites SJER and LENO were moved to avoid overlap with other charts and a red arrow indicates the true location of the site. **b)** Principal coordinates (PC) analysis shows how samples group by vegetation class based on their soil, climate, and vegetation factors. One point represents one soil sample (*n* = 1199 soil samples). Soil abiotic and vegetation variables (blue), climate variables (magenta), and microbial variables (black) are shown. Parentheses on the axis labels indicate the amount of variation explained by each PC axis. Abbreviations include nitrogen (N), ectomycorrhizal fungi (EMF), ectomycorrhizal (EM), mean annual temperature (MAT) and mean annual precipitation (MAP). **c)** Spearman correlations show associations between net ammonification or nitrification rates and the relative abundances of microbial groups, soil, climate, and vegetation factors across the entire dataset (bacterial variables *n* = 805 soil samples, fungal variables *n* = 927, climatic variables and soil inorganic N *n* = 4675, soil moisture *n* = 4512, soil C:N ratio *n* = 935, soil temperature *n* = 4645, soil pH *n* = 4496, percent EM tree basal area *n* = 4667). A microbial group is shown if it was included in our hypotheses or met the criteria outlined in the Methods: Statistical Analysis.

## RESULTS

### Opposite microbial taxonomic and functional groups correlate with ammonification and nitrification rates

We found novel associations between microbial functional and taxonomic groups and net ammonification or nitrification rates in soils across the United States. Across the entire dataset, the strongest positive correlates of net ammonification rates were the relative abundances of the bacterial classes Acidobacteriae and Bacteroidia, the fungal class Saccharomycetes, and EMF (Fig. 1c, Supplementary Fig. 1). Meanwhile, the strongest positive correlates of net nitrification were the relative abundances of the bacterial class Thermoleophilia and pathogenic fungi (Fig. 1c, Supplementary Fig. 1). Latitude and longitude explained variation in both rates (Supplementary Table 2), but microbial groups explained a significant amount of variation when accounting for spatial autocorrelation across samples (Supplementary Table 3). Opposite variables correlated with ammonification versus nitrification rates across the dataset (Fig. 1c), even though the two rates were not significantly negatively correlated across the dataset (ρ = −0.01, P = 0.50). Our finding that completely different groups of microorganisms are associated with net ammonification versus nitrification rates reinforces the concept that soil ammonification and nitrification processes are fundamentally unique, driven by different organisms and environmental conditions, and should be represented as such in biogeochemical models.

Microbial groups known for their N-decomposition abilities correlated positively with net ammonification rates across the dataset: as we hypothesized, EMF correlated positively with net ammonification, but oligotrophic bacteria as a whole did not (Fig. 1c). EMF can produce N-decomposing enzymes in soils that release ammonium from organic matter^23–25^, potentially driving ammonification, but they can also thrive in conditions that promote ammonification by other microbes, such as high soil moisture and low pH^12^. Random forest analysis showed that the top 11 taxa whose abundances drove the relationship between EMF and net ammonification rates all have contact or short-distance hyphal exploration types (Supplementary Fig. 2a), which are hypothesized to degrade more organic N than medium- and long-distance exploration types^26^. Though net ammonification was not correlated with oligotrophic bacteria as a group, it was correlated with bacteria in the class Acidobacteriae (Fig. 1c), which are oligotrophs with the genomic capacity to degrade complex N-containing polysaccharides^27–29^. Bacteroidia, which also correlated positively with net ammonification rates, are neither oligotrophs nor copiotrophs, but could promote ammonification through protein degradation and the production of N-targeting CAZymes^30,31^. Indeed, the top Bacteroidia taxa accounting for the positive relationship with net ammonification were often from the family Chitinophagaceae, which degrade chitin, a major N-source in soil^32^ (Supplementary Fig. 2c). Similarly, Saccharomycetes, a geographically widespread class of yeasts, are free-living decomposers^33^ that can produce abundant extracellular enzymes, including N-decomposing enzymes like chitinases^34^.

In line with our hypothesis, net nitrification rates correlated positively with the relative abundance of copiotrophic bacteria as a group, as well as the copiotrophic bacterial classes Thermoleophilia and Gemmatimonadetes and plant and animal pathogenic fungi (Fig. 1c). Copiotrophic bacteria, Thermoleophilia, and Gemmatimonadetes^35–38^ all contain denitrifying bacteria, which use the products of nitrification to produce gaseous nitrogen compounds like the greenhouse gas nitrous oxide and dinitrogen gas^20^. Random forest analysis showed that the most important taxa driving the relationships between net nitrification and copiotrophic bacteria, Gemmatimonadetes, or Thermoleophilia are all denitrifying bacteria or uncharacterized bacteria (Supplementary Fig. 3a-c)^35,36,39^. Microbial nitrification activity likely promotes growth of denitrifying (nitrous oxide reducing) bacteria by increasing soil nitrous oxide concentrations^40^, potentially explaining the positive correlations between net nitrification rates and microbial groups containing denitrifying bacteria. In addition to denitrifying bacteria, copiotrophic bacteria include many nitrifying bacteria^7,8^, but interestingly, copiotrophic bacteria correlated more strongly positively with net nitrification than nitrifying bacteria alone (Fig. 1c), suggesting that the copiotroph group could contain previously uncharacterized nitrifying bacteria or non-nitrifying copiotrophs are increasing their growth in response to high nitrification rates. We also found that fungal pathogens of plants and animals were positively associated with nitrification (Fig. 1c). Because pathogenic fungi tend to be nitrophilic^21^ and are more diverse in high N soils globally^12^, pathogens may not be performing nitrification, but rather increasing in abundance when nitrification rates are high.

### Each microbe-N cycling relationship occurs under specific environmental conditions

Because microbial functional groups can also track climate or other environmental changes^41^, we explored whether the microbial groups associated with net ammonification or nitrification rates represented particular environmental conditions where these rates were highest. We found that microbial groups correlating positively with net ammonification (Acidobacteriae, Bacteroidia, Saccharomycetes, and EMF) were associated with environmental conditions common in forested ecosystems (especially deciduous forests) and higher latitudes: high soil moisture and percent ectomycorrhizal (EM) trees, but low mean annual temperature, mean diurnal temperature range, soil temperature, and soil pH (Fig. 1b, Supplementary Fig. 4). In contrast, most microbial groups that correlated positively with net nitrification rates (Gemmatimonadetes, Thermoleophilia, and fungal pathogens) were associated with environmental conditions characteristic of shrubland ecosystems, grasslands, and croplands: high soil pH and mean diurnal range, but low soil moisture, percent of EM trees, and mean annual precipitation (Fig. 1b, Supplementary Fig. 4), most frequently found in the central United States (Fig. 1a). Nevertheless, the environmental conditions where a microbial group was most abundant were not always where its relationship with N cycling rates was the strongest.

### Net ammonification

We found that every relationship between a microbial group and net ammonification was present under a specific set of environmental conditions (Fig. 2). Acidobacteriae (which are oligotrophs) were significantly associated with net ammonification rates across more environmental conditions than any other microbial group and across diverse ecosystem types (Fig. 2a), indicating that this class is a key widespread determinant of net ammonification rates. The relative abundance of Acidobacteriae explained the most variation in net ammonification rates in shrublands (29% of variation), soils with high concentrations of inorganic N (19%), non-EM tree dominated ecosystems (14%), during winter-to-spring seasonal transitions (12%), and in climates with high mean diurnal temperature range (12%, Fig. 2a, Supplementary Table 4). 84% of shrubland samples in the dataset are non-EM dominated (< 50% EM basal area) and have high mean diurnal range (>14°C, Fig. 2b), representing a specific sub-ecosystem where Acidobacteriae may drive ammonification (Fig. 2c). Shrublands encompass a wide geographic range in our dataset, from Alaska to the southwestern and central United States (Fig. 1a). Acidobacteriae are widely geographically distributed and can be core members of shrubland soil bacterial communities^42,43^. Shrublands and high inorganic N may promote the N-decomposition activity of Acidobacteriae, which is consistent with previous work showing that bacteria, rather than fungi, can dominate decomposition in non-forest ecosystems^44^ and high soil N suppresses fungi more so than bacteria^5^.

**Fig. 2.**
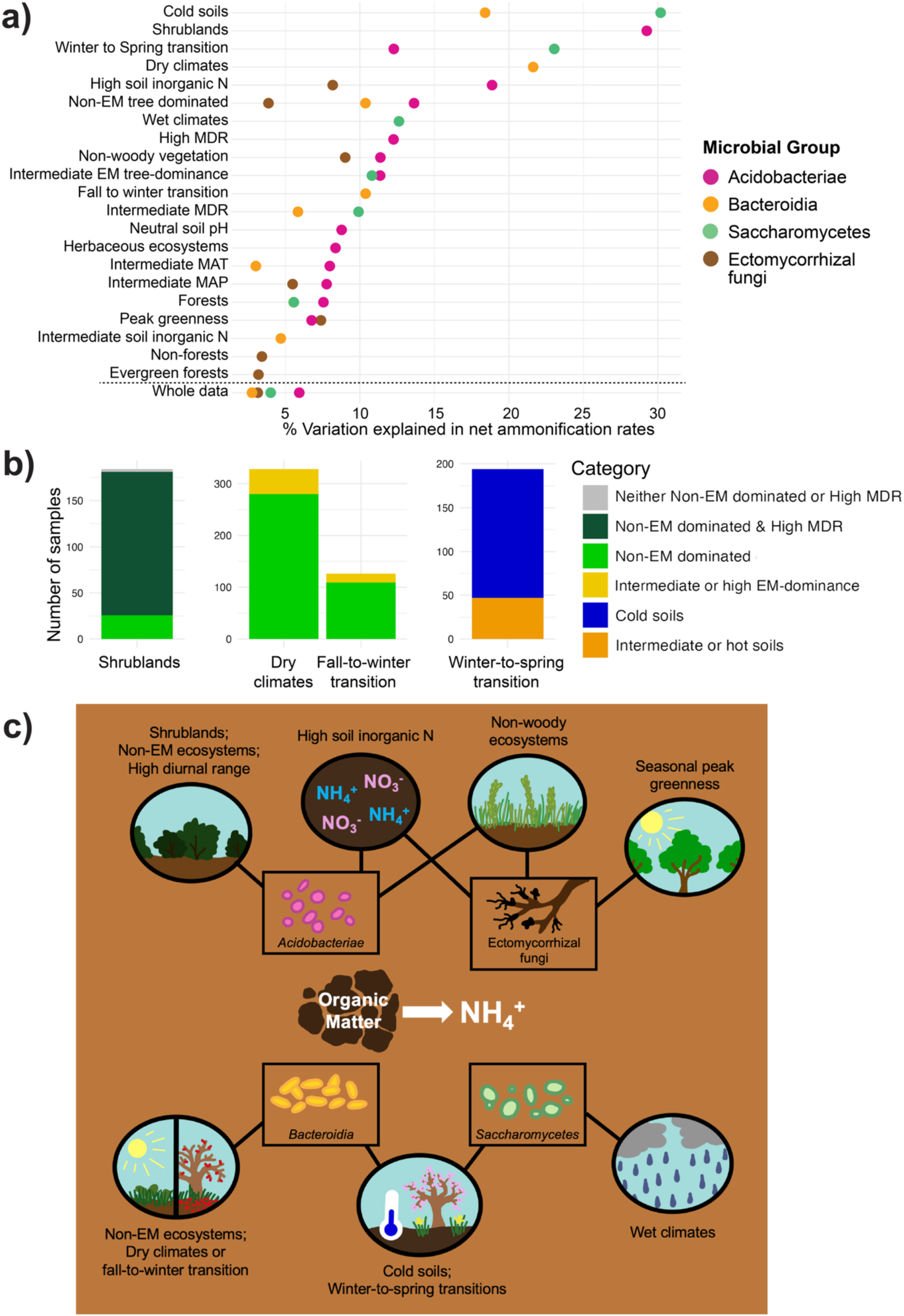
Environmental conditions control the relationships between microbial functional groups and net ammonification rates. **a)** The relative abundances of Acidobacteriae, Bacteroidia, Saccharomycetes, and ectomycorrhizal fungi significantly explain net ammonification rates under the environmental conditions listed on the Y-axis. The X-axis shows the percent variation in net ammonification rates explained by a microbial group in each environmental condition listed on the Y-axis. One point represents one model. Model results are only shown if the relationship is significant (P < 0.001) and the model’s R^2^ is greater than the R^2^ of the model using the whole dataset. Detailed statistical outputs for each model and sample sizes for each data subset are shown in Supplementary Tables 1 and 4. **b)** Bar plots showing the number of samples belonging to overlapping environmental conditions in which microbial groups explain the most variation in net ammonification rates. **c)** Summary diagram showing the environmental conditions in which microbial functional and taxonomic groups explain the most variation in net ammonification rates. Black boxes indicate microbial groups, while black ovals indicate environmental conditions. A microbial group is connected to an environmental condition with a black line if the microbial group explains high variation in net ammonification rates in that environmental condition. Abbreviations include nitrogen (N), ectomycorrhizal (EM), mean diurnal temperature range (MDR), mean annual temperature (MAT) and mean annual precipitation (MAP).

The non-oligotrophic bacterial class Bacteroidia explained variation in net ammonification rates primarily in dry climates (22% of the variation), cold soils (18%), non-EM dominated ecosystems (10%), and during fall to winter seasonal transitions (10%, Fig. 2, Supplementary Table 4). Dry climates and cold temperatures typically reduce microbial decomposition activity^45^, but Bacteroidia are known to utilize diverse carbon sources, produce many organic matter-decomposing enzymes^31^, and remain active in cold conditions^46,47^. 85% of samples from dry climates (mean annual precipitation <800 mm) and 87% of samples collected during fall-to-winter seasonal transitions are also from non-EM dominated ecosystems (Fig. 2b), representing a unique environment where Bacteroidia are associated with ammonification (Fig. 2c).

The relative abundance of Saccharomycetes, a common genus of yeasts, explained the most variation in net ammonification rates in cold soils (30% of the variation), during winter-to-spring transitions (23%), in wet climates (13%), intermediate EM-dominance ecosystems (11%), and climates with moderate daily temperature variability (10%, Fig. 2, Supplementary Table 4). Cold soils (< 10°C soil temperature) span 12 sites and 10 vegetation cover types, including forests, shrublands, wetlands, grasslands, and croplands. Because 76% of samples collected during winter-to-spring transitions had cold soils (Fig. 2b), Saccharomycetes may be associated with net ammonification during winter-to-spring transitions because of colder soil temperatures during this season (Fig. 2c). Yeasts can often withstand cold conditions^48^ and have physiological adaptations that allow them to continue growth at freezing temperatures^49^, which may allow Saccharomycetes to drive or respond to ammonification in cold environments.

The relative abundance of EMF explained variation in net ammonification rates in ecosystems dominated by non-woody vegetation (9% of the variation), soils with high inorganic N (8%), during seasonal peak greenness (7%), and in moderate precipitation climates (5%, Fig. 2a, Supplementary Table 4). None of these conditions had large overlap (Fig. 1b, Supplementary Fig. 5c), instead representing distinct environments in which EMF are positively associated with net ammonification rates (Fig. 2c). These patterns may be partially driven by short-distance exploration type EMF, which were positively associated with net ammonification rates under these same conditions (Supplementary Table 4), and helped explain the relationship with net ammonification rates across the entire dataset (Supplementary Fig. 2a). The positive association in ecosystems dominated by non-woody vegetation is surprising, but many of the non-woody ecosystems where EMF were present in our dataset often had one or two EM plants present. In this case, EMF may exert an outsized impact on localized N cycling when they are present, but not dominant, perhaps because of reduced competition with other EMF^50^. EMF explained more variation than other microbial groups during peak greenness (Fig. 2a,c), potentially due to increased plant carbon investment belowground when EMF are most active^51,52^.

### Net nitrification

Many relationships between microbial groups and net nitrification rates were found in wet, hot, and high-N environments. In support of our hypothesis, the relative abundance of pathogenic fungi explained the highest amounts of variation in net nitrification rates in wet climates (19% of the variation), evergreen forests (14%), and soils with intermediate moisture (11%, Fig. 3, Supplementary Table 4). None of these environmental conditions had large overlap (Fig. 1c, Supplementary Fig. 5e), representing unique conditions in which pathogenic fungi are associated with net nitrification. Pathogens may be utilizing the abundant nitrate produced from nitrification for their own growth in these environments^21,53^, or their plant and animal hosts may be more abundant and diverse where soil nitrate is high^54,55^.

**Fig. 3.**
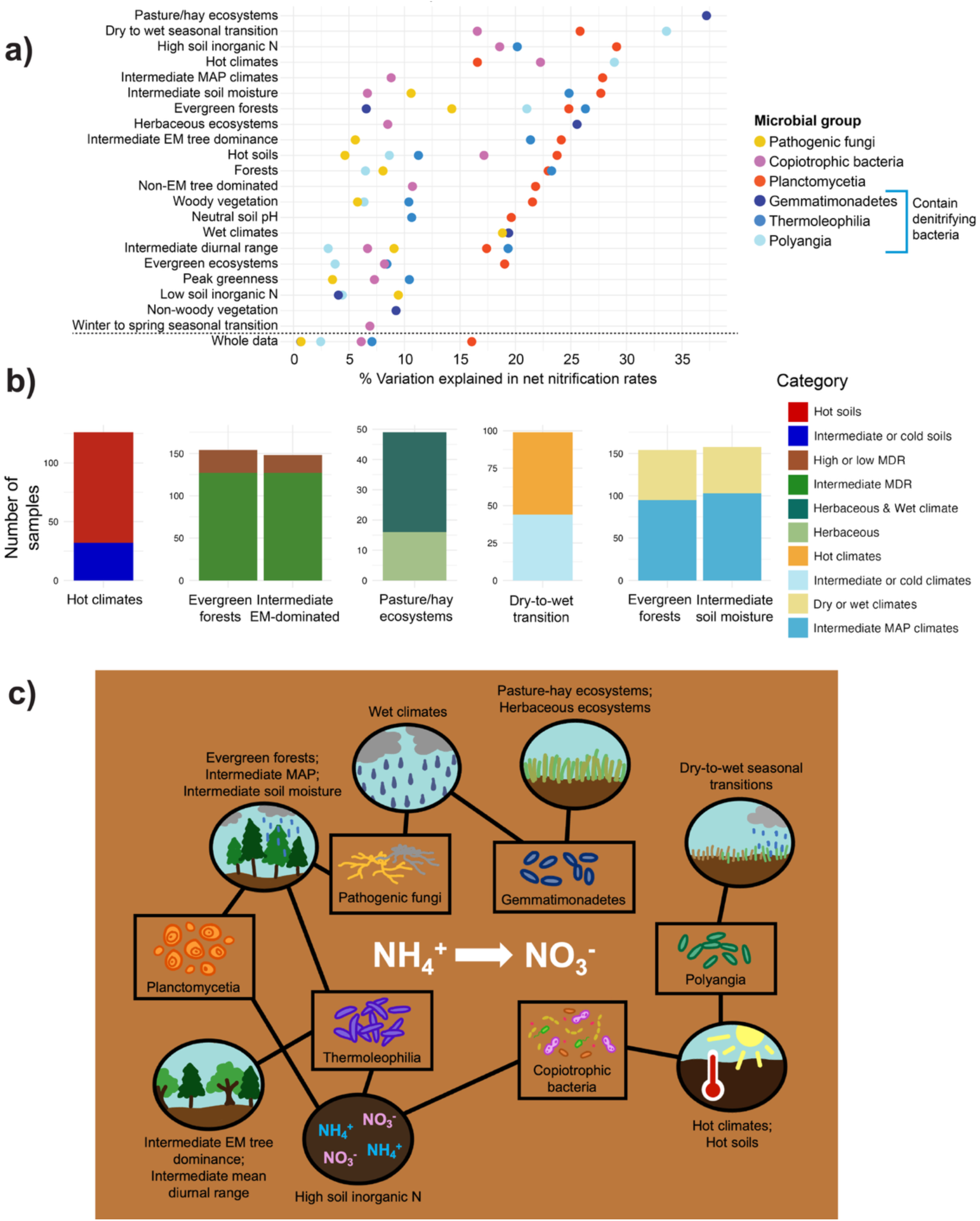
Environmental conditions control the relationships between microbial functional groups and net nitrification rates. **a)** The relative abundances of plant and animal pathogenic fungi, copiotrophic bacteria, and bacterial classes significantly explain net ammonification rates under the environmental conditions listed on the Y-axis. The X-axis shows the percent variation in net ammonification rates explained by microbial groups in each environmental condition listed on the Y-axis. One point represents one model. Model results are only shown if the relationship is significant (P < 0.001) and the model’s R^2^ is greater than the R^2^ of the model using the whole dataset. Detailed statistical outputs for each model and sample sizes for each data subset are shown in Supplementary Tables 1 and 4. **b)** Bar plot showing the number of samples belonging to overlapping environmental conditions in which microbial groups explain the most variation in net nitrification rates. **c)** Summary diagram showing the environmental conditions in which microbial functional and taxonomic groups explain the most variation in net nitrification rates. Black boxes indicate microbial groups, while black ovals indicate environmental conditions. A microbial group is connected to an environmental condition with a black line if the microbial group explains high variation in net nitrification rates in that environmental condition. Abbreviations include nitrogen (N), ectomycorrhizal (EM), mean diurnal temperature range (MDR), and mean annual precipitation (MAP).

Also in support of our hypothesis, we found that copiotrophic bacteria explained variation in net nitrification rates in hot climates (22% of the variation), soils with high inorganic N (19%), high soil temperatures (17%), and during dry-to-wet seasonal transitions (17%, Fig. 3, Supplementary Table 4). 75% of samples from hot climates (mean annual temperature >18°C) also had high soil temperatures (> 20°C, Fig. 3b). Previous work shows that copiotrophic bacteria are abundant at forest edges, which are characterized by high soil temperatures and high inorganic N^17^, conditions that could increase nitrification activity by nitrifier copiotrophs^56^, which in turn could potentially promote growth and activity of other non-nitrifying bacteria within the copiotroph group (e.g., denitrifying bacteria). Thermoleophilia and Gemmatimonadetes, copiotrophic classes of bacteria that contain denitrifying bacteria^35–38^, explained variation in net nitrification rates across different environmental conditions than the broader group of copiotrophic bacteria (Fig. 3). Thermoleophilia explained the most variation in net nitrification rates in evergreen forests (26% of the variation), soils with intermediate moisture (25%), intermediate-EM dominated ecosystems (21%), soils with high inorganic N concentrations (20%), and climates with intermediate mean diurnal range (19%, Supplementary Table 4). 86% of intermediate-EM dominated ecosystems (50 - 90% EM trees) and 82% of evergreen forests had intermediate mean diurnal temperature ranges (12 - 14°C, Fig. 3b), which indicates a specific sub-ecosystem where Thermoleophilia is associated with nitrification (Fig. 3c). Meanwhile, Gemmatimonadetes explained the most variation in net nitrification rates in pasture/hay ecosystems (37% of the variation), herbaceous ecosystems (26%), and climates with high mean annual precipitation (19%, Supplementary Table 4). All pasture/hay samples were herbaceous, and 67% of pasture/hay samples were from wet climates (mean annual precipitation > 1100 mm, Fig. 3b), suggesting a strong link between Gemmatimonadetes and nitrification within herbaceous, wet ecosystems (Fig. 3c) despite the wide distribution across latitudes, from the southernmost site in the contiguous U.S. to the mid-Atlantic region (Fig. 1a). Pastures and wet environments often have high denitrification rates, which can promote the buildup of nitrous oxide in soil^57^, promoting metabolism of nitrous oxide-reducing bacteria within Gemmatimonadetes. Interestingly, other non-copiotrophic bacterial classes containing denitrifying bacteria were also associated with net nitrification in wet, warm conditions: for example, the class Polyangia positively explained more variation in net nitrification rates than any other microbial group during dry-to-wet seasonal transitions (34%) and in hot climates (29%, Fig. 3, Supplementary Table 4). Many of the top taxa responsible for the association between Polyangia and net nitrification were from the genus *Haliangium* (Supplementary Fig. 3e), which can express multiple genes involved in denitrification^58^.

Unexpectedly, the relative abundance of Planctomycetia, a class of oligotrophic bacteria, explained variation in net nitrification rates in more environmental conditions than any other microbial group, including 16% of the variation across the entire dataset (Fig. 3a). While there are no known nitrifying bacteria within Planctomycetia, the phylum Planctomycetes (which contains Planctomycetia) includes anaerobic ammonia-oxidizing bacteria^59^. Few terrestrial Planctomycetia have been characterized^60^, so it is possible that some uncharacterized bacteria within Planctomycetia perform ammonia oxidation. For example, the top Planctomycetia taxa with the strongest relationships to net nitrification are all from unknown species (Supplementary Fig. 3f). Planctomycetia explained the most variation in net nitrification rates in soils with high inorganic N concentrations (29% of the variation), intermediate mean annual precipitation (28%), intermediate soil moisture (28%), during dry-to-wet seasonal transitions (26%), and in evergreen forests (25%, Fig. 3, Supplementary Table 4). 65% of soils with intermediate moisture (30-50% water content) and 62% of samples from evergreen forests are from moderately wet climates (mean annual precipitation 800 - 1100 mm, Fig. 3b). Moderately wet environments and high inorganic N are necessary for anaerobic ammonia oxidation^61–63^. Planctomycetia also explained more variation in net nitrification rates than any other microbial group in soils with neutral pH (Fig. 3a), which is notable because neutral to basic pH promotes nitrification activity by bacteria but not of archaea^3^.

## DISCUSSION

In a world where natural nutrient cycles are increasingly disturbed by global change factors like warming climates, increasing atmospheric N deposition, and fertilizer use, it is critical to understand the fundamental processes controlling the availability of key soil nutrients to achieve a sustainable global future^64^. In this study, we provide a novel foundational understanding of microbe-N cycling relationships across the United States. We identified previously-unknown associations between N cycling rates and microbial groups, showing that these associations are dependent on vegetation cover, soil, and climate variables. Often, the environmental conditions under which a given microbe-N cycling relationship was strongest are consistent with conditions known to promote the N-decomposition, nitrification, or denitrification activity of the microbial group.

A major outcome from this work is confirmation that ammonification and nitrification should be treated as separate biogeochemical and ecological processes, associated with entirely different groups of microorganisms and environmental conditions (Fig. 1c). While our results provide critical information needed to explicitly include soil microbes into biogeochemical cycling models^6^, we suggest that ammonification and nitrification should not be merged into a singular “net N mineralization” measure of soil N cycling processes, as is done in the majority of ecological soil N-cycling studies and models^65–67^. Modeling ammonium versus nitrate fluxes separately may also be advantageous because these two N forms differentially affect plant N uptake^68–70^ and soil nitrate can contribute to waterway eutrophication^71^. A key step to explicitly incorporate microbial functional groups into large-scale biogeochemical models will be to use microbial group abundances to estimate net ammonification and nitrification rates, alongside abiotic and plant variables, under conditions where those relationships were found to exist. This approach is increasingly feasible, as bacterial and fungal amplicon sequence datasets are becoming more widespread and publicly accessible across the world^12,72^.

Our findings generate novel hypotheses about the ecology of widespread soil microbial functional and taxonomic groups. Acidobacteriae, Bacteroidia, Saccharomycetes, and EMF, for example, may contribute to ammonification in diverse environments through production of N-decomposing enzymes^25,28,30^. Categorizing bacteria into copiotrophs can aid in distilling the diversity of microbial communities into a broad group that can explain and predict nitrification rates in warm, N-rich conditions, while Planctomycetia may be an especially widespread group driving or responding to nitrification rates across many environments. Multiple bacterial classes containing denitrifiers (Thermoleophilia, Gemmatimonadetes, Polyangia) are closely associated with net nitrification in environmental conditions known to promote denitrification (e.g., pastures, N-rich soils, hot and wet environments, Fig. 3), suggesting an interaction between nitrification, denitrifiers, denitrification, and environmental factors. These results can aid efforts to predict abundances of denitrifying microbes and potentially rates of nitrous oxide (a powerful greenhouse gas) production and consumption. All of the bacterial and fungal classes mentioned here have many undescribed species^33,60,73^, highlighting the need for more studies characterizing the functions of species within these groups to complete our understanding of soil N cycling. Overall, our work defining the context-dependency of microbial-N cycling relationships is a critical step towards linking microbial community composition to function in diverse ecosystems.

## METHODS

### NEON sampling design

NEON is a coordinated ecological monitoring network across the United States and its territories, including terrestrial sampling sites that cover 20 eco-climatic domains^74^. The NEON terrestrial sampling design is described in detail in Hinckley et al. (2016)^75^, Barnett et al. (2019)^76^, and Weintraub-Leff et al. (2023)^22^. At each of the 19 sites used in this study, soil sampling was performed in ten 40 x 40 m plots, in which sampling locations are identified by stratified random sampling. These plots cover the major vegetation and land-cover types (according to the National Land Cover Database) within each site. During a given sampling timepoint, three soil cores were collected from each plot and split into mineral and organic horizons when possible. During a sampling year, soil samples are collected at three timepoints: seasonal peak greenness and the seasonal transitions before and after peak greenness. We only included samples in our analyses that had a complete set of microbial, inorganic N cycling, soil abiotic, and vegetation data associated with the sample. Each soil sample used for analysis was associated with unique microbial and soil abiotic data, with no repeated measurements on the same samples. To ensure the same sampling and processing protocols were used for all samples (i.e., avoid changes in protocols over time), we only used samples collected from 2017 - 2018. This approach resulted in 805 samples with associated bacterial community data and 927 samples with fungal community data. These samples spanned 19 sites (16 in the continental U.S., 2 in Alaska, 1 in Puerto Rico) with data from 8-10 plots at each site, encompassing 10 vegetation cover classes (Fig. 1a). The number of plots at each site, sampling dates, number of samples per site, and data availability at each site is summarized in Supplementary Table 5.

### Microbial composition

We utilized NEON’s existing soil bacterial and fungal community composition data to determine the relative abundance of different microbial functional and taxonomic groups, which was measured via amplicon sequencing of the 16S v3-4 and ITS1 rDNA regions. DNA extraction protocols, primers, and sequencing protocols are described in detail in NEON TOS Protocol and Procedure: Soil Biogeochemical and Microbial Sampling (NEON.DOC.014048vL)^77^, NEON DNA Extraction Standard Operating Procedure v.1^78^, and Qin et al (2021)^79^. Briefly, microbial DNA was extracted from 0.25 g of soil using the PowerSoil HTP Kit (QIAGEN, Hilden, Germany). Amplicons were sequenced using an Illumina Miseq with v2 2×250 base-pair paired end chemistry at Argonne National Laboratory and Illumina MiSeq with v3 2×300 base-pair paired end chemistry at Battelle Applied Genomics. Raw sequences can be accessed via the NEON data portal (data.neonscience.org)^80^.

Processing of the microbial sequence data, including bioinformatics, quality filtering, and taxonomy and functional group assignment is described in Werbin et al. (2024)^7^. Briefly, bioinformatic processing and quality filtering of the sequence data was completed following the dada2 pipeline^81^ to identify Exact Sequence Variants (ESVs). Subsequent quality filtering steps, percent read retention, and pipeline parameters are described in Qin et al. 2021^79^. Bacterial and fungal samples had a median of 10,935 (range 5,025 - 105,819) and 17,730 (range 5,013 - 262,419) sequences, respectively, after filtering steps and removing low-quality samples. To account for the compositional nature of microbial amplicon sequencing data, all sequence abundances were converted to relative abundances by dividing each ASV count in each sample by the total number of reads in that sample, which is an appropriate normalization method when regressing the relative abundances of microbial taxa against separate, non-compositional response variables (e.g., N cycling rates)^82,83^. Taxonomy was assigned to ESVs with the rAnomaly pipeline using both the UNITE (v8.2) and UTOPIA (90, 2019-08) databases for fungi and the SILVA v138 and GTDB v95 databases for bacteria. 78% of bacterial ESVs and 35% of fungal ESVs were able to be assigned taxonomy at the class rank. 40% of bacterial ESVs and 18% of fungal ESVs were able to be assigned taxonomy at the genus rank.

In this study, we chose to focus on microbial functional groups that are abundant across ecosystems at large spatial scales, including groups that were previously found to be associated with ammonification or nitrification rates at single sites^10,17^: ectomycorrhizal fungi, saprotrophic fungi, plant and animal pathogenic fungi, copiotrophic bacteria, and oligotrophic bacteria. Fungal functional groups were assigned at the genus rank using the FUNGuild and Fungal Traits databases^84,85^. To test for bacterial copiotroph and oligotroph group relationships with process rates, we used an in-house database compiled from literature reviews and genomic pathway presence to assign bacterial ESVs as copiotrophs, oligotrophs, or neither based on their taxonomy at the genus through phylum ranks^7,8,16,86^. This database is hosted on Github at https://github.com/zoey-rw/soil_bacteria_functional_groups. To capture microbial functions associated with distinct taxonomic groups, we also ran analyses using the top 20 most abundant classes of bacteria and fungi (Supplementary Table 6). We chose to group at the class level because many bacterial N-cycling functions and fungal ecological lifestyles are conserved at the class level^87–89^ and microbial classes are often widespread across large geographical ranges^90^. To assess whether specific EMF exploration types were responsible for EMF-N cycling relationships, we also ran analyses using EMF exploration types.

### Soil abiotic data

Soil net N mineralization and nitrification rate data were measured by NEON staff using the “buried bag” method^22^. Net rates of N mineralization and nitrification were obtained using the ‘neonNTrans’ package (v 0.4)^91,92^. To correct for contamination in sample blanks, we used the “Option 4” dataset that includes blank-corrected inorganic N values, which allows for the largest dataset yet avoids zero-inflation of the data^22^. To calculate net ammonification rates, we subtracted net nitrification rates from net N mineralization rates. Protocols for net N mineralization and nitrification measurements are described in detail in Weintraub-Leff et al. (2023)^22^. Microbial and soil physicochemical data were collected on the same samples used for the initial ammonium and nitrate measurements. Soil temperature data were obtained from NEON DP1.00041.001^93^, soil water content (i.e., soil moisture) data was obtained from NEON DP1.00094.001^94^, and soil pH and C:N content were obtained from NEON DP1.10086.001^95^.

### Vegetation data

The vegetation cover type of each sampling plot was classified by NEON using the National Land Cover Database (NLCD)^96^. Vegetation cover types spanned the classes “cultivated crops,” “deciduous forest,” “dwarf scrub,” “evergreen forest,” “grassland herbaceous,” “mixed forest,” “pasture hay,” “sedge herbaceous,” “shrub scrub,” and “woody wetlands”. We used NLCD class to separate plots based on ecosystem type, deciduous versus evergreen, and woody versus non-woody vegetation-dominated as described in Supplementary Table 1. To quantify the dominance of EM-associated plants, which are known to affect soil N cycling and fungal communities^97^, we calculated the proportion of EM trees in each plot using stem diameter values from the NEON woody plant vegetation structure dataset (DP1.10098.001)^98^ as described in Werbin et al. (2024)^7^.

### Climate data

We downloaded mean annual temperature, mean annual precipitation, and mean diurnal temperature range (with higher values indicating larger daily temperature fluctuations) data associated with each sampling plot from the WorldClim database^99^. We extracted these climatic variables associated with the spatial coordinates of each sampling plot using the ‘geodata’ (v 0.6.6)^100^ and ‘terra’ (v 1.8.70)^101^ R packages.

### Statistical analysis

All statistical analyses were run in R version 4.4.2^102^. To understand how soil, climate, and vegetation variables were grouped across the dataset, we ran a Principal Component Analysis (PCA) using the ‘princomp’ function in the ‘stats’ package^102^. We used net ammonification rates, net nitrification rates, soil temperature, soil moisture, soil pH, mean annual temperature, mean annual precipitation, mean diurnal temperature range, and the proportion of EM trees as input variables to the PCA. To visualize how microbial functional and taxonomic groups are associated with these environmental factors, we mapped vectors for each microbial group on top of the PCA using the ‘envfit’ function in the ‘vegan’ package (v 2.6.8)^103^. To determine how these continuous environmental variables varied with vegetation class, we ran Kruskal-Wallis tests using the ‘FSA’ (v 0.10.0)^104^ and ‘multcompView’ (v 0.1.10)^105^ packages.

Because abiotic and vegetation variables within and across sites were often structured by latitude and longitude (Fig. 1a-b), our analyses accounted for latitude, longitude, and spatial autocorrelation in microbial communities. To assess whether associations between microbial functional groups and net ammonification or nitrification rates were due to spatial autocorrelation, we ran multiple regression models on distance matrices of net ammonification or nitrification rates as the response variable, and the microbial group and spatial distance between samples as the predictor variables^17^ using the ‘lm’ function in the ‘stats’ package (v 4.4.2)^102^. Then, to test for correlations between microbial abundances, net ammonification or nitrification rates, and environmental variables across the whole dataset, we ran Spearman correlations between all soil, microbial, vegetation, and climate variables using the ‘Hmisc’ package (v 5.2.1)^106^, with significance levels adjusted for multiple corrections to P < 0.001 with the Bonferroni method^107^. To assess which taxa accounted for the relationship between a microbial group and the N cycling rate, we ran random forest models with either net ammonification or nitrification as the response variable and the relative abundances of individual taxa (ESVs) as the predictors. We chose which taxa to include as predictors in each random forest model by selecting all taxa that had a positive and significant (P < 0.05) Pearson correlation with the response variable. Random forest models were trained on 70% of the input data and tested on the other 30% of the input dataset, and final models were run using the ‘randomForest’ function in the ‘randomForest’ package (v 4.7.1.1)^108^ with n trees = 2000.

To determine the environmental context dependencies of relationships between microbial groups and net ammonification or nitrification rates, we constructed generalized linear models in each environmental condition data subset (Supplementary Tables 1 & 4) using the ‘glm’ function in the ‘stats’ package (v 4.4.2) with the relative abundance of a microbial group as the explanatory variable and either net ammonification or nitrification rates as the response variable. To account for spatial effects on N cycling rates, we included latitude and longitude as additional predictors within each model rather than site, because site is confounded with ecosystem/vegetation type (i.e., different sites have different vegetation types), many sites are clustered together in space (Fig. 1a), and much of the between- and within-site variation in abiotic and climate factors varies with latitude and longitude. We also ran separate generalized linear models explaining net ammonification and nitrification rates using only latitude and longitude as predictors. Based on the distribution of the response variables, we set the family parameter to Gamma with link = log in all models and added a pseudocount of 10 to net ammonification and 12 to net nitrification rates to turn all values positive. We calculated the pseudo R^2^ of each model using McFadden’s R^2^ ^109^. To calculate percent variance explained by the microbial predictor, we used the R^2^ of the model including only the microbial group as a predictor (without latitude and longitude), because including latitude and longitude often increased the R^2^ of the model, reflecting additional variance explained by geographic location (Supplementary Table 4). Each model was tested for overdispersion using the ‘testDispersion’ function in the ‘DHARMa’ package (v 0.4.6)^110^. We ran each model on the entire dataset, then ran the same models on data subsetted by vegetation cover type (NLCD class), sample timing (season), and high/medium/low EM tree dominance, soil inorganic N content, soil pH, soil moisture, soil temperature, soil C:N, mean annual precipitation, mean annual temperature, and mean diurnal range. When binning continuous variables into high/medium/low categories, thresholds were determined based on environmental and biological relevance (for example, known soil pH ranges between different forest types and microbial pH tolerances) and the distribution of values across the dataset (Supplementary Table 1).

To visualize the microbial groups that explained the most variation in net ammonification and nitrification rates, we focused on the microbial functional groups in our hypotheses, as well as a subset of microbial classes that explained the most variation in N cycle process rates. A class was selected for data visualization if it met three criteria: 1) it was significant within the model in 5 or more environmental conditions (with a Bonferroni-corrected P-value of P < 0.001 when latitude and longitude were also included as predictors); 2) it had a positive association with net ammonification or nitrification rates; and 3) it explained more than 10% of the variance in net ammonification or 15% of the variance in net nitrification rates in 3 or more environmental conditions. For models run on subsetted data, we selected models where the R^2^ was greater than the R^2^ of the model run on the entire dataset. For every selected model, if it appeared that a relationship was driven by 3 or fewer samples, those samples were removed. If the model no longer met our criteria for selection after removal of these points, the model was not used.

## ACKNOWLEDGEMENTS

This material is based in part upon work supported by the National Ecological Observatory Network (NEON), a program sponsored by the U.S. National Science Foundation (NSF) and operated under cooperative agreement by Battelle. We would like to thank the NEON staff for their work collecting the data used in this study and making it publicly accessible. This work was supported by U.S. Department of Energy Biological and Environmental Research awards DE-SC0020403 and DE-SC0012704 and the Ecotechnology and Landscape Restoration Research Fund provided by Leap of Faith Technologies, Inc. to Jennifer M. Bhatnagar and an NSF Graduate Research Fellowship (10.13039/100023581, solicitation 19-590) and American Association of University Women American Dissertation Fellowship to Corinne Vietorisz.

## DATA AVAILABILITY STATEMENT

All data used in this study are publicly accessible via the NEON data portal (data.neonscience.org). The specific datasets used are listed in the methods with full citations, DOIs, and web links to each dataset located in the References^80,93–95,98^.

