## Supplementary_Information for "Soil nitrogen cycling rates are linked to microbial functional and taxonomic groups across the United States"

### *Supplementary Tables*

Supplementary Tables are located in the “Supplementary_Tables” file.

**Supplementary Table 1. Data subsets input to generalized linear models based on categories of plant, soil, and climate variables.**

**Supplementary Table 2. Statistical outputs from generalized linear models using latitude and longitude to explain net ammonification and nitrification rates.**

**Supplementary Table 3. Statistical outputs from multiple regression models on distance matrices.** All models had the formula: lm(dist(Response variable) ~ dist(Microbial predictor) + dist(spatial coordinates).

**Supplementary Table 4. Statistical outputs from generalized linear models run on data subsets.**

**Supplementary Table 5. Sampling dates, number of plots, number of samples, and data availability by sampling site.**

**Supplementary Table 6. Top 20 most abundant fungal and bacterial classes tested for relationships with net ammonification and nitrification rates.**

### *Supplementary Figures*

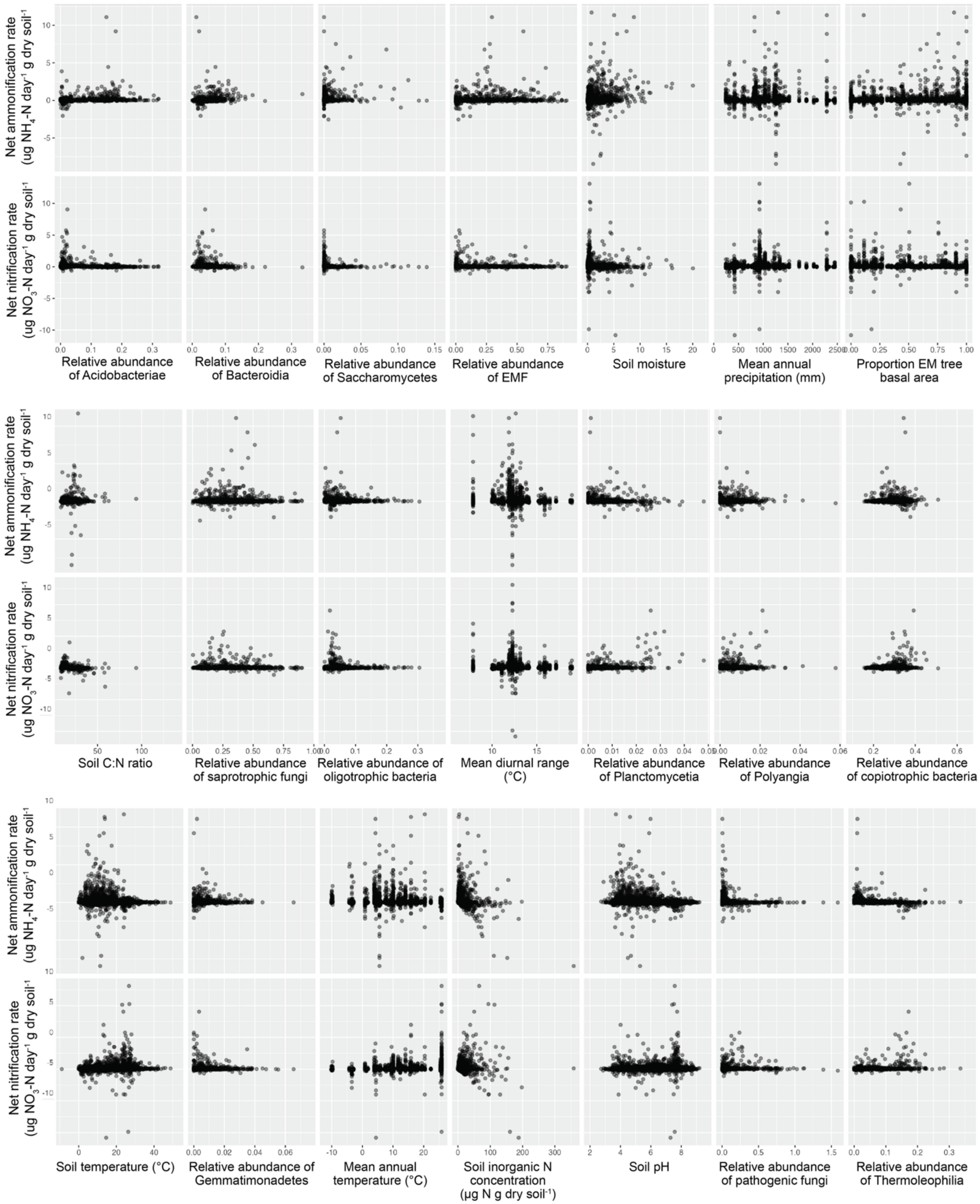

**Supplementary Figure 1. Scatterplots showing the relationships between net ammonification or nitrification rates and microbial, soil, climate, and vegetation factors across the entire dataset.** Abbreviations include ectomycorrhizal fungi (EMF) and ectomycorrhizal (EM).

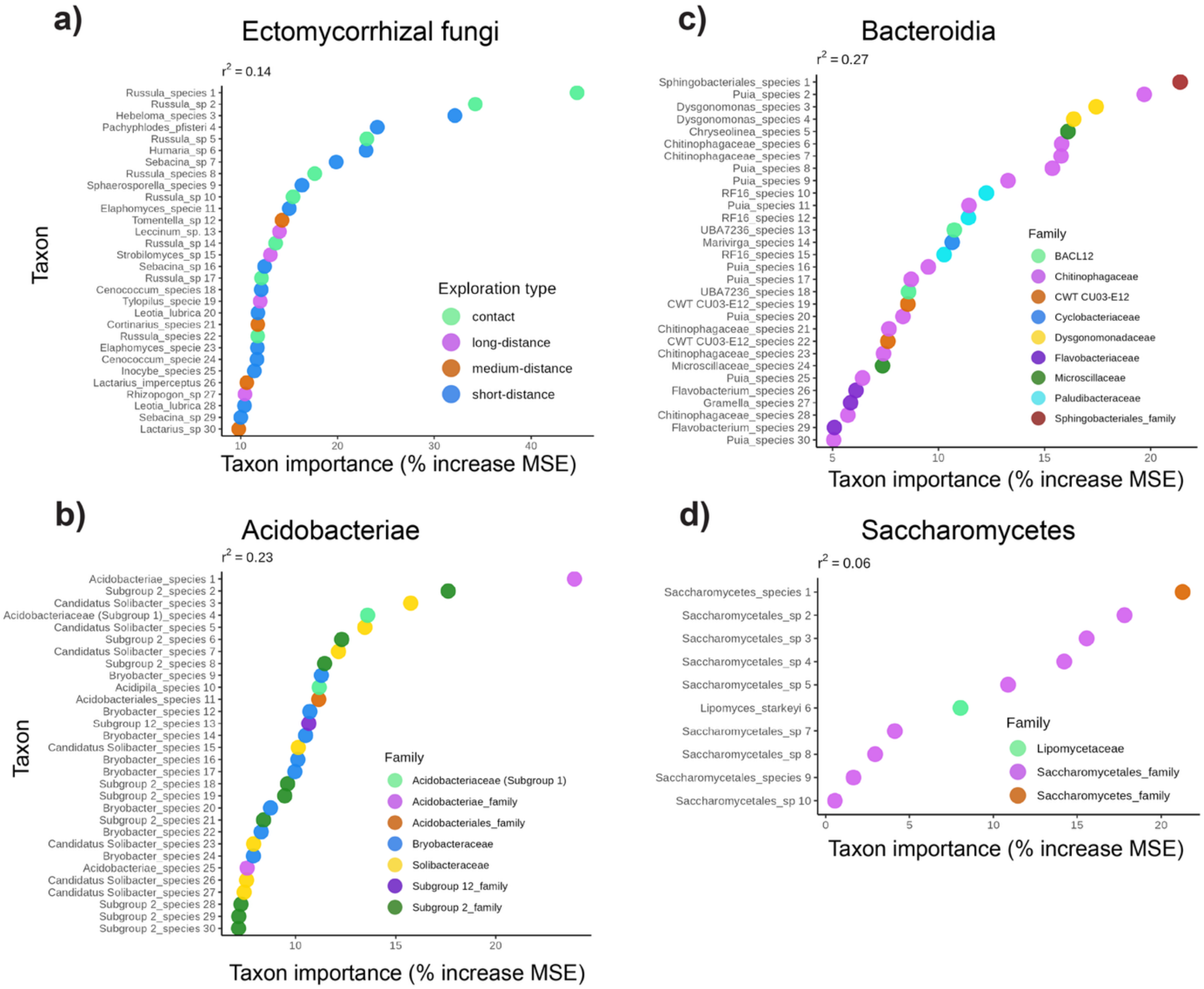


**Supplementary Figure 2. Top 30 taxa of a) ectomycorrhizal fungi, b) Acidobacteriae, c) Bacteroidia, and d) Saccharomycetes driving each group's positive relationship with net ammonification.** To assess which taxa drove the relationship between a microbial group and net ammonification, we ran random forest models with net ammonification as the response variable, and the relative abundances of individual taxa (ESVs) as the predictors. One panel represents one random forest model. Individual taxa used as predictors in the random forest model are shown on the Y axes. X axes show the importance of each taxon within the random forest model, where a higher % increase in mean square error (MSE) indicates that the taxon plays a larger role in driving the relationship with net ammonification. One point represents one taxon.

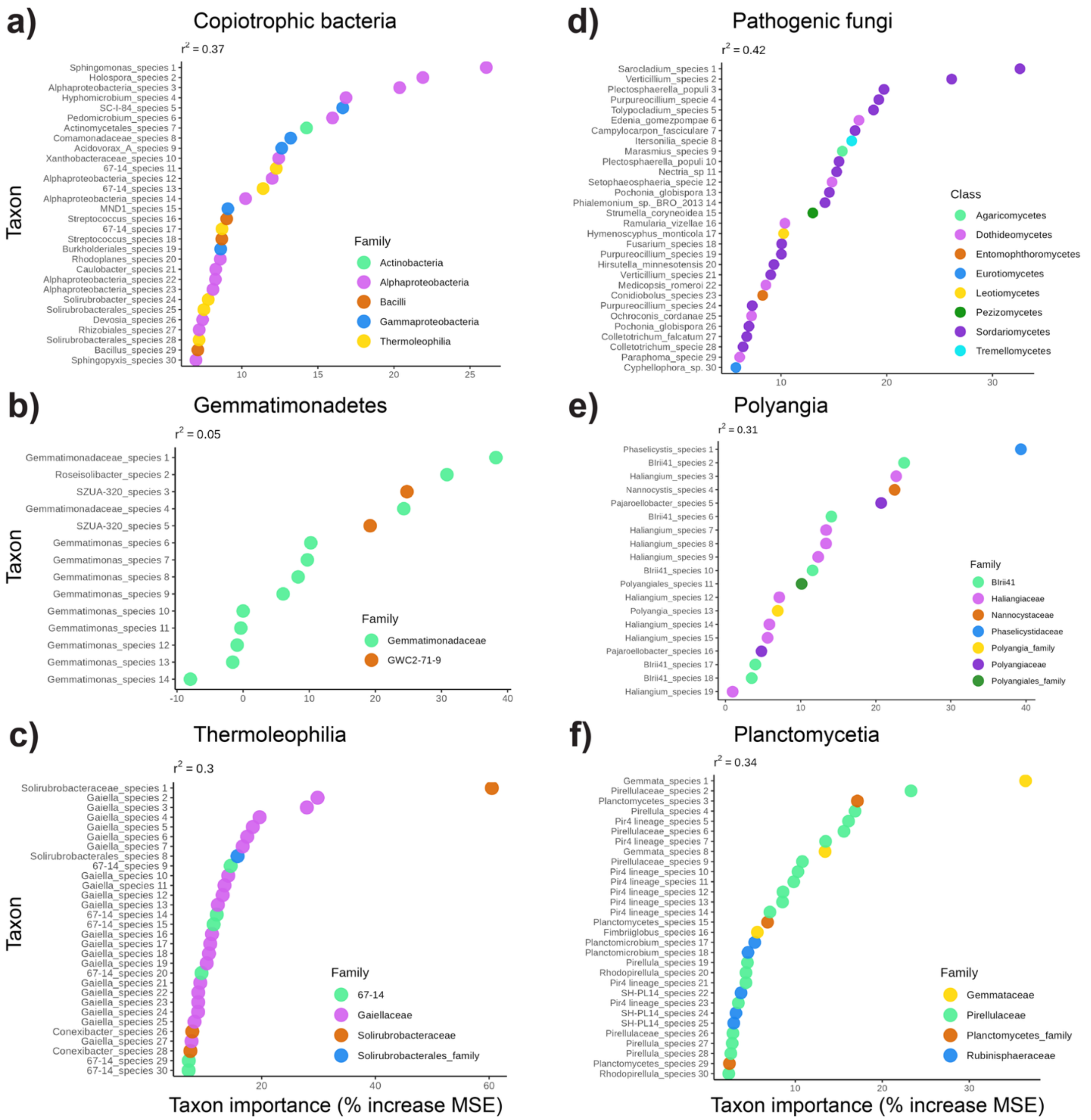


**Supplementary Figure 3. Top 30 taxa of a) copiotrophic bacteria, b) Gemmatimonadetes, c) Thermoleophilia, d) pathogenic fungi, e) Polyangia, and f) Planctomycetia driving each group's positive relationship with net nitrification.** To assess which taxa drove the relationship between a microbial group and net nitrification, we ran random forest models with net nitrification as the response variable, and the relative abundances of individual taxa (ESVs) as the predictors. One panel represents one random forest model. Individual taxa used as predictors in the random forest model are shown on the Y axes. X axes show the importance of each taxon within the random forest model, where a higher % increase in mean square error (MSE) indicates that the taxon plays a larger role in driving the relationship with net nitrification. One point represents one taxon.

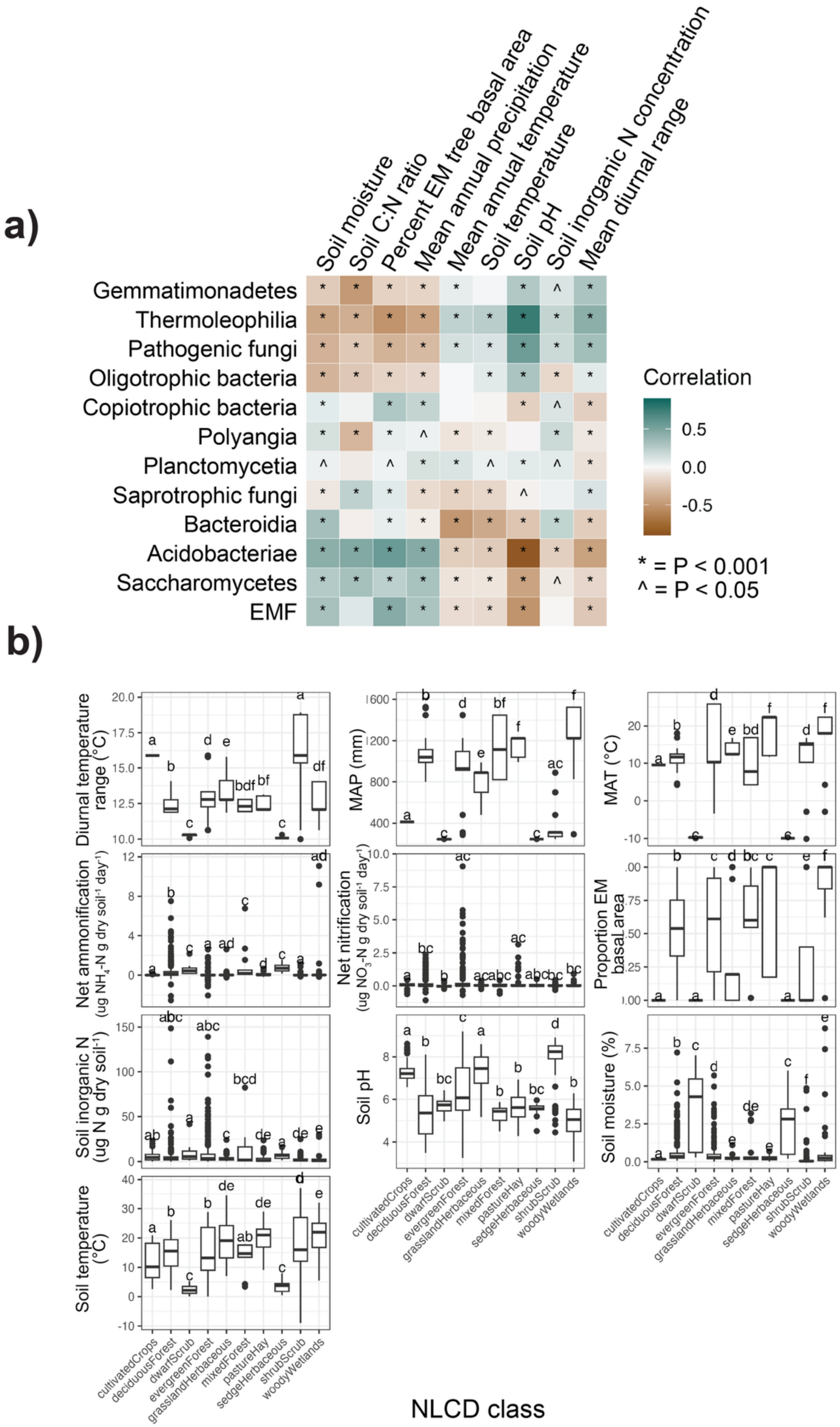
a)
Soil moisture
Soil C:N ratio
Percent EM tree basal area
Mean annual precipitation
Mean annual temperature
Soil temperature
Soil pH
Soil inorganic N concentration
Mean diurnal range
Gemmatimonadetes
Thermoleophilia
Pathogenic fungi
Oligotrophic bacteria
Copiotrophic bacteria
Polyangia
Planctomycetia
Saprotrophic fungi
Bacteroidia
Acidobacteriae
Saccharomycetes
EMF
Correlation
0.5
0.0
-0.5
* = P < 0.001
^ = P < 0.05
b)
Diurnal temperature range (°C)
MAP (mm)
MAT (°C)
Net ammonification (ug NH4-N g dry soil-1 day-1)
Net nitrification (ug NO3-N g dry soil-1 day-1)
Proportion EM basal area
Soil inorganic N (ug N g dry soil-1)
Soil pH
Soil moisture (%)
Soil temperature (°C)
cultivatedCrops
deciduousForest
dwarfScrub
evergreenForest
grasslandHerbaceous
mixedForest
pastureHay
sedgeHerbaceous
shrubScrub
woodyWetlands
NLCD class

**Supplementary Figure 4. Climate, soil, and vegetation variables correlate with microbial groups and vary with National Land Cover Database (NLCD) vegetation class. a)** Spearman correlations show associations between the relative abundances of microbial groups and soil, climate, and vegetation factors. Abbreviations include ectomycorrhizal fungi (EMF) and ectomycorrhizal (EM). **b)** Letters above each boxplot denote statistically significant differences between vegetation classes as determined by pairwise Kruskal-Wallis tests. Boxes represent the interquartile range of the data, midpoint lines represent the median, whiskers represent data within 1.5 times the interquartile range, and points represent outliers. Abbreviations include mean annual precipitation (MAP), mean annual temperature (MAT), and ectomycorrhizal (EM).

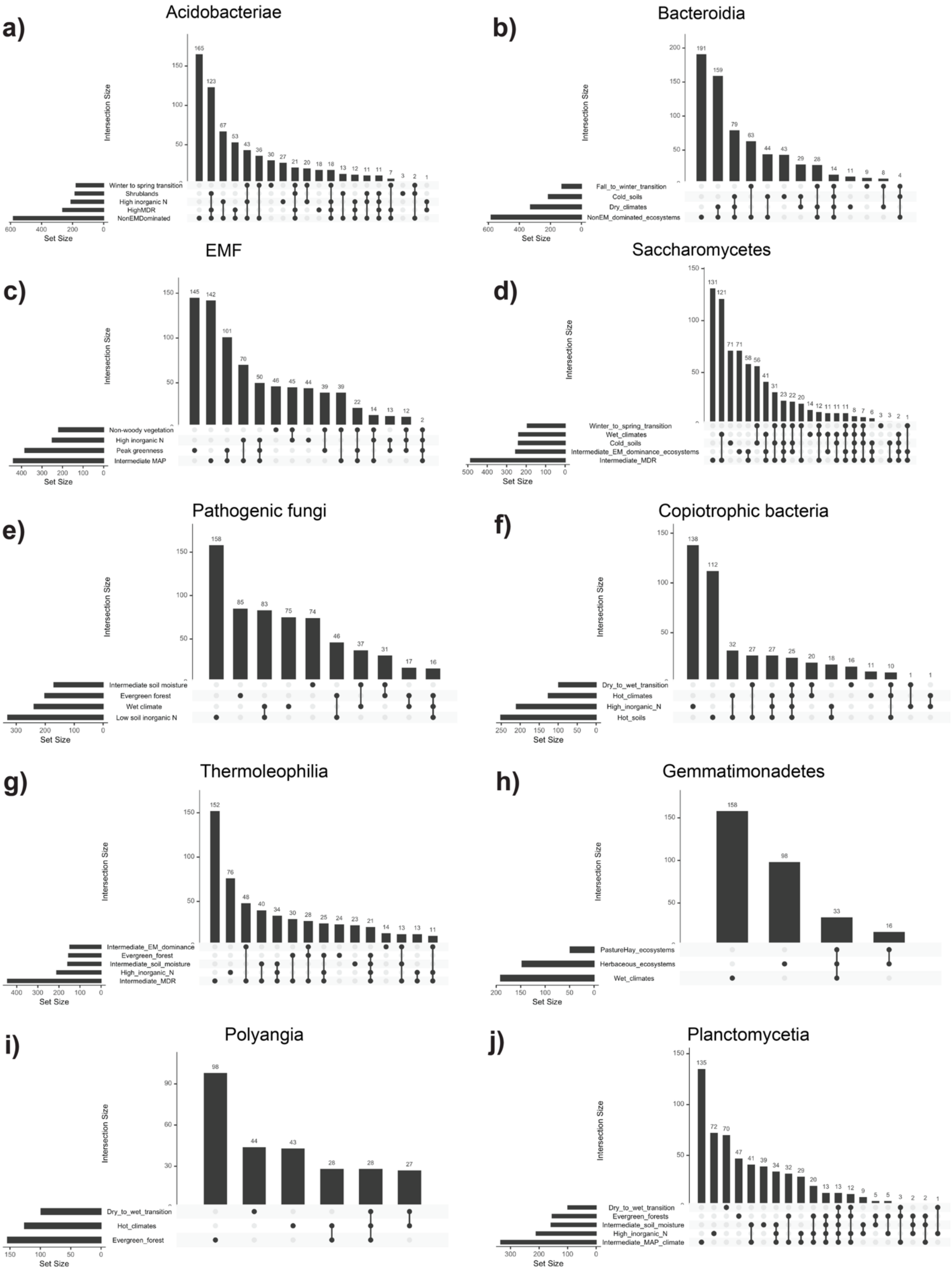
a)
Acidobacteriae
Intersection Size
165
123
67
53
43
36
30
27
21
20
18
18
13
12
11
11
7
3
2
1
Winter to spring transition
Shrublands
High inorganic N
HighMDR
NonEMDominated
Set Size
b)
Bacteroidia
Intersection Size
191
159
79
63
44
43
29
28
14
11
9
8
4
Fall_to_winter_transition
Cold_soils
Dry_climates
NonEM_dominated_ecosystems
Set Size
c)
EMF
Intersection Size
145
142
101
70
50
46
45
44
39
39
22
14
13
12
2
Non-woody vegetation
High inorganic N
Peak greenness
Intermediate MAP
Set Size
d)
Saccharomycetes
Intersection Size
131
121
71
71
58
56
41
31
23
22
20
14
12
11
11
11
8
7
6
3
3
2
1
Winter_to_spring_transition
Wet_climates
Cold_soils
Intermediate_EM_dominance_ecosystems
Intermediate_MDR
Set Size
e)
Pathogenic fungi
Intersection Size
158
85
83
75
74
46
37
31
17
16
Intermediate soil moisture
Evergreen forest
Wet climate
Low soil inorganic N
Set Size
f)
Copiotrophic bacteria
Intersection Size
138
112
32
27
27
25
20
18
16
11
10
1
1
Dry_to_wet_transition
Hot_climates
High_inorganic_N
Hot_soils
Set Size
g)
Thermoleophilia
Intersection Size
152
76
48
40
34
30
28
25
24
23
21
14
13
13
11
Intermediate_EM_dominance
Evergreen_forest
Intermediate_soil_moisture
High_inorganic_N
Intermediate_MDR
Set Size
h)
Gemmatimonadetes
Intersection Size
158
98
33
16
PastureHay_ecosystems
Herbaceous_ecosystems
Wet_climates
Set Size
i)
Polyangia
Intersection Size
98
44
43
28
28
27
Dry_to_wet_transition
Hot_climates
Evergreen_forest
Set Size
j)
Planctomycetia
Intersection Size
135
72
70
47
41
39
34
32
29
20
13
13
12
9
5
5
3
2
2
1
Dry_to_wet_transition
Evergreen_forests
Intermediate_soil_moisture
High_inorganic_N
Intermediate_MAP_climate
Set Size

**Supplementary Figure 5. UpSet plots showing intersections between the top environmental conditions in which a) Acidobacteriae, b) Bacteroidia, c) ectomycorrhizal fungi (EMF), and d) Saccharomycetes explain the most variance in net ammonification rates, and in which e) plant and animal pathogenic fungi, f) copiotrophic bacteria, g) Thermoleophilia, h) Gemmatimonadetes, i) Polyangia, and j) Planctomycetia explain the most variance in net nitrification rates.** Horizontal bars on the left indicate the total sample size of each data subset. The matrix of connected dots identifies which sets are included in each intersection: filled dots represent membership in that intersection, and connected lines link all sets involved. Vertical bars above the matrix show the number of samples belonging to each corresponding intersection. Abbreviations include mean annual precipitation (MAP), mean diurnal temperature range (MDR), and ectomycorrhizal (EM).
